# Late embryonic expansion of a novel bone ridge underlies the evolutionary transformation of cylindrically shaped forelimb bones into the flattened skeleton of the penguin flipper

**DOI:** 10.64898/2026.06.29.735166

**Authors:** Charlie Longtine, Hannah A. Grunwald, Stephen Treaster, Matthew P. Harris, Clifford J. Tabin

## Abstract

The evolution of flippers for wing-powered diving in penguins is a striking example of tetrapod limb specialization. The modern penguin flipper is structurally reinforced by a characteristic dorsoventral ‘flattening’ of the long bones accompanied by a reduction in distal forelimb musculature, features which emerged convergently in flightless diving birds and aquatic mammals. While an extensive fossil record informs the morphological sequence through which these changes occurred, the evolutionary pressures and developmental mechanisms underlying these modifications are unknown. We find that in avian and mammalian forelimbs, a flattened bone morphology only emerged in aquatic lineages that lost ancestral modes of locomotion, including in flightless diving birds, pinnipeds, and cetaceans. Using penguin embryos as an accessible model for investigating flipper development, we demonstrate that early patterning of forelimb musculoskeletal morphology is similar to that seen in forelimbs of non-aquatic birds.

Instead, later modifications of gene expression and cell and tissue behaviors underlie flipper phenotypes. Thus, we find that in the early penguin forelimb, the initial cues that pattern the muscle do not differ from other avian species, however late embryonic changes in proliferation result in dramatic reduction of muscle. Likewise, forelimb bones in penguins initially have similar cross-sectional proportions to those in flighted birds. The shape of these bones is, however, remodeled late in embryonic development through a process that shares molecular hallmarks with bone ridge formation at tendon attachment sites. In these bones, ridge-forming tissue initiates at the ends of the bones (the epiphyses) and extends into tendon-like connective tissue along the lateral edges of the bone, widening the long bones along the anterior-posterior axis and producing a ‘flattened’ bone. Using spatial transcriptomics and comparative genomic tools we determine that differentially expressed genes between the ridge-forming tissues and long bone cartilage are significantly enriched for signals of selection in the penguin lineage and that these genes may also be convergently evolving in marine mammals. Together, these data show that the evolution of musculoskeletal morphology in the penguin flipper occurred through expansion or novel deployment of molecular programs typically associated with tendon-attachment sites during late embryonic development.

## Introduction

The developmental mechanisms through which individual bones get their distinctive shapes during development and across vertebrate evolution remain largely unknown. The initial pattern of long bones in tetrapod limbs is formed by condensations of chondrogenic limb bud mesenchymal cells, which establish the location and general shape of the future skeletal element. These condensations are initially similar in size[1,2] and the differences in the lengths of various limb skeletal elements emerge through differential proliferation rates[3], differences in the extent of hypertrophy of chondrocytes[4–7], and timing of growth cessation[8]. In addition to precisely proportioned length, individual long bones have distinctive shapes due to localized bony outgrowths that form ridges, providing stable anchoring sites for connective tissues such as tendons and ligaments[9].

Dramatic shifts in the morphology of skeletal elements have occurred between species over the course of evolution. Modifications of bone proportion and shape, particularly in the limb, have allowed for innovations in vertebrate locomotion which ultimately facilitate expansion into otherwise inaccessible niches. For example, birds and bats convergently evolved modified skeletal elements in the forelimb that permitted powered flight, while caecilians, skinks, and snakes convergently evolved dramatic limb reduction to facilitate lateral undulation locomotion[10].

One particularly interesting example of a derived limb modification is the vertebrate flipper, which arose independently in turtles, penguins, cetaceans (whales and dolphins), and pinnipeds (seals and sea lions). All of these lineages underwent secondary marine transitions, in which tetrapods that were adapted for life on land gave rise to aquatic descendants. In each case, these animals evolved specialized forelimbs characterized by a flattened shape with apparent lateral broadening or dorsoventral thinning of the limb that facilitates efficient underwater locomotion. While only some of these groups use their flippers directly for propulsion, all make use of the distinct morphology of the flipper as a hydrodynamic ‘foil’ to maximize the efficiency of locomotion through water[11,12].

Notably, within birds, wing-propelled diving has independently evolved multiple times[13], however, among the extant wing-propelled divers - eg. penguins, diving petrels (*Pelecanoides)*, shearwaters (*Puffinus*), gannets (*Morus*), and dippers (*Cinclidae*) - only penguins have evolved a flipper-like forelimb morphology. Penguins are one of the most specialized groups of extant birds, having evolved a variety of unique adaptations to their marine environment. Among the suite of morphological, physiological, and behavioral adaptations in penguins are numerous modifications of forelimb structure, including tendonization and loss of distal musculature[14–16], an extreme flattening of the forelimb bones, shortening of the wing, and changes in the proportional length of wing segments[17],[18]. Together, these modifications allow the penguin flipper to act as a stiff paddle and hydrofoil during sustained wing-propelled diving[11,12].

While penguins represent the only extant group of flipper-bearing birds, a number of distantly related flightless diving birds with flipper-like morphologies are seen in the avian fossil record, including plotopterids, mancalline or Lucas auks, and the great auk. Morphological similarities in the wing skeletal structure between these independent extant and extinct lineages of flightless diving birds highlight convergent adaptive evolution in the Aves forelimb skeleton to increase efficiency of wing-propelled diving, though differences in muscular anatomy indicate that the morphological convergence of the musculature is limited[19].

The observed convergence in morphology among these groups raises the possibility of parallelism in the processes that underlie the development of this morphology. However, it is unknown if there are shared developmental mechanisms among vertebrate groups in forming the specialized structures of the flipper. To get a handle on this question, we turned our attention to the development of the penguin flipper. The close evolutionary relationship between penguins and the related taxa that do not share their dramatic morphological forelimb adaptations, coupled with the ability to obtain eggs for investigating embryonic development, make the penguin a powerful case study to understand how changes in development lead to adaptive change of the skeleton in flipper-bearing vertebrates.

Here, we compare forelimb long bone morphology among diving mammals and birds and show parallel evolutionary transitions that result in flattened forelimb bones in these lineages. To understand the cellular and molecular mechanisms that contribute to proportion and shape of the musculoskeletal structures in the flipper, we then interrogate the development of the penguin forelimb. We demonstrate that the modifications of penguin forelimb bones are not established through changes in initial bone patterning. Instead, the shift in the cross-sectional proportion, or aspect ratio, of these bones is achieved through expansion of epiphyseal bone ridges along the anterior and posterior edges of the forelimb bones late during embryonic development. These bone ridges co-express tendon and cartilage markers, making them highly reminiscent of bone ridges that form at tendon attachment sites in both adaptive and aberrant bony protuberances. Remarkably, the extension of bone ridges along the length of the bones during penguin embryogenesis mirrors the evolutionary trajectory of altered aspect ratio in extinct stem penguins. Coincident with the expansion of the bone ridge, musculature in the distal penguin flipper is progressively reduced in the distal penguin forelimb as a result of a reduction in muscle proliferation. Spatial transcriptomics of the bony ridge reveal differentially expressed genes that are enriched for genomic selection in the penguin lineage and may be convergently evolving in marine mammal transitions. Collectively, these data support a model wherein the penguin lineage evolved the highly derived flipper by progressively modifying late-stage limb development.

## Results

### Forelimb skeletal flattening emerges convergently in aquatic birds and mammals that lost alternate locomotion

We first aimed to characterize evolutionary patterns of bone flattening in birds. Distinct changes within the forelimb anatomy of diving avian lineages have been previously documented, however, the degree of skeletal change across the avian phylogeny has not been described[18]. To quantify forelimb bone morphology and flattening, we measured the height (dorsal-ventral) and width (anterior-posterior) of the radius, ulna, and tibia diaphyses. We used these values to calculate the aspect ratio (height/width) of each skeletal element across a broad taxonomic sampling of birds, using ulnar and radial values together for each species to compare forelimb bones as a whole between clades (Fig 1B-D and S1A-C). Aspect ratios close to 1.0 indicate a rounded shape, whereas aspect ratios < 1.0 are flattened in the dorsoventral plane and those > 1.0 are flattened in the lateral (anterior-posterior) plane.

**Fig 1.**
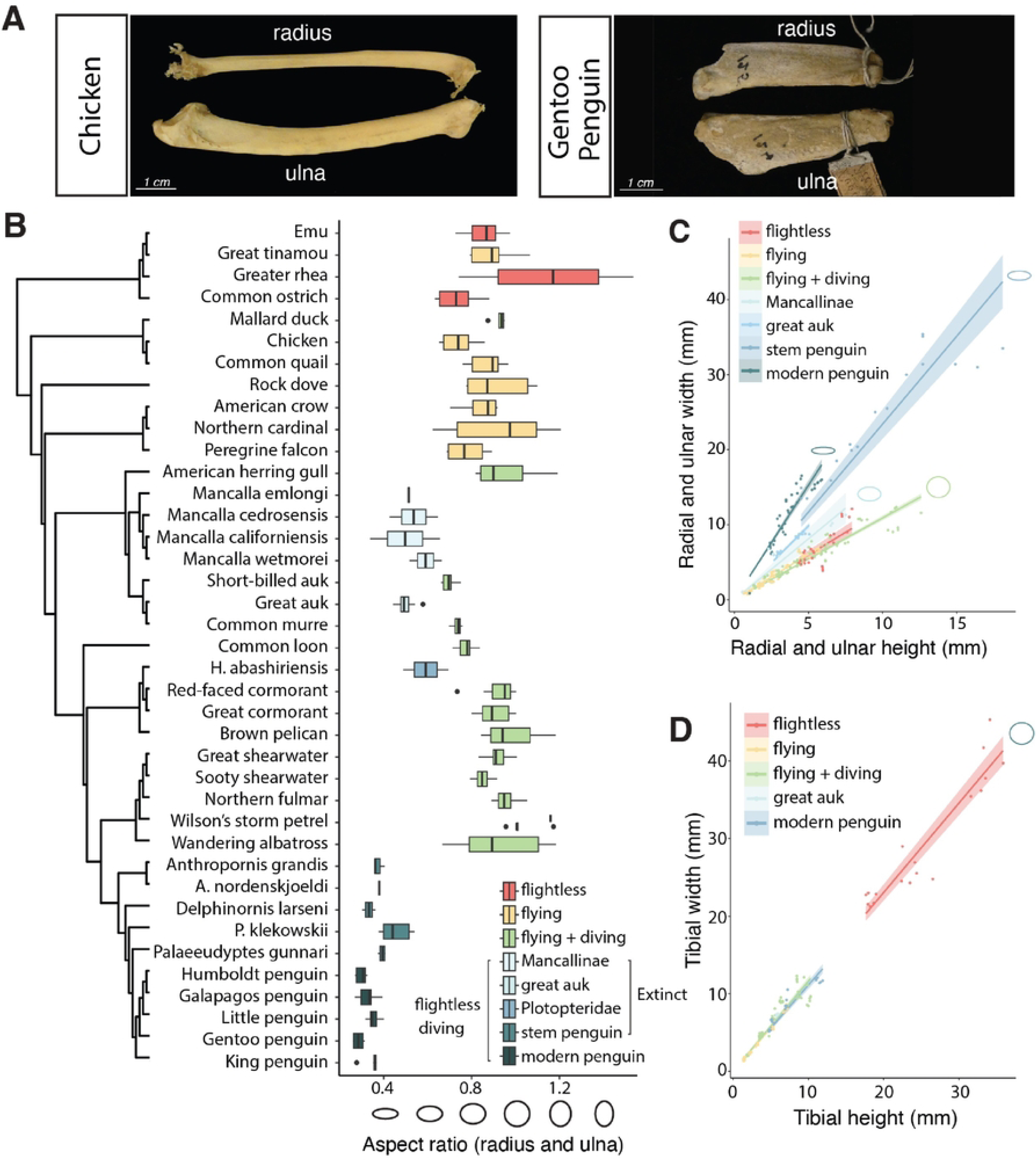
Forelimb bone flattening convergently emerges in flightless diving birds. (A) Chicken (*Gallus gallus)* and gentoo penguin (*Pygoscelis papua)* radius and ulna (B) Mean cross-sectional aspect ratio (AR) of the radius and ulna for extant and extinct avian species. Ovals below the graph represent bone cross-sections at points along the x-axis. AR = 1.0 is round. AR < 1.0 is dorso-ventrally flattened. (C, D) Regression of bone height vs width by avian sub-group for radius/ulna (C) and tibia (D). The slope of each line reflects the aspect ratio. Ovals represent the bone cross-section for different lines.

Mean combined radial and ulnar aspect ratio did not significantly differ between flighted birds, flightless paleognaths, and a broad phylogenetic distribution of flighted diving birds (adj. p-value > 0.9880 all comparisons). In contrast, radial and ulnar aspect ratio was significantly lower in penguins compared to each of these groups, indicating significantly flatter forelimb bones (adj. p-value < 0.0001 all comparisons) (Fig S1C). Tibial aspect ratio was consistent across all phyla measured, indicating that flat bones are a forelimb-specific adaptation in penguins, rather than a general characteristic of penguin bone development (Fig 1D).

To determine whether extinct stem penguins shared the modifications to the radius and ulna that we observed in modern penguins, we took advantage of the rich literature describing fossil penguin specimens. Penguins have an extensive fossil record[20–23] dating the origin of penguins to ∼60 million years ago (MYA) with the crown penguin group dating to only ∼15 MYA. Using photographs and measurements published in the literature, we calculated the aspect ratio of forelimb bones from fossils of extinct penguin species. Within stem penguins, mean aspect ratios of radius and ulna were significantly lower than extant flightless, flighted, and flighted diving birds (adj. p-value < 0.0001 all comparisons). An extension of this analysis to independent lineages of extinct flightless diving birds showed similar trends, with Mancallinae[24] showing significantly flatter bones compared to extant non-penguins (adj. p-value vs flighted and flighted diving < 0.0001; flightless = 0.0001) (Fig 1B-C and S1C). Although the mean aspect ratios in the extinct flightless diving plotopterid, *H. abashiriensis*[25], and the great auk were notably lower than that seen in flighted birds, only a single species was available from these clades so significance was not tested. Modern penguins exhibited the most extreme skeletal morphology of any avian group measured, with the lowest mean aspect ratio, however this difference was only modestly significantly different when compared with Mancallinae (adj. p-value = 0.0228) and not significant when compared with stem penguins (adj. p-value = 0.8886) (Fig S1C).

These results demonstrate that dorso-ventral flattening of forelimb bones evolved independently in at least three avian lineages, exclusively in lineages that were both flightless as well as diving. The repeated shifts in forelimb bone aspect ratio within birds highly adapted to navigating an aquatic environment raised the possibility that similar morphologies might be observed in aquatic species outside Aves, reflecting a broader pattern of adaptation or constraint. To determine whether other vertebrate divers shared a similar degree of forelimb bone flattening, we measured the aspect ratio of the radius and ulna in aquatic mammals and related outgroups (Fig 2, S2, and S3).

**Fig 2.**
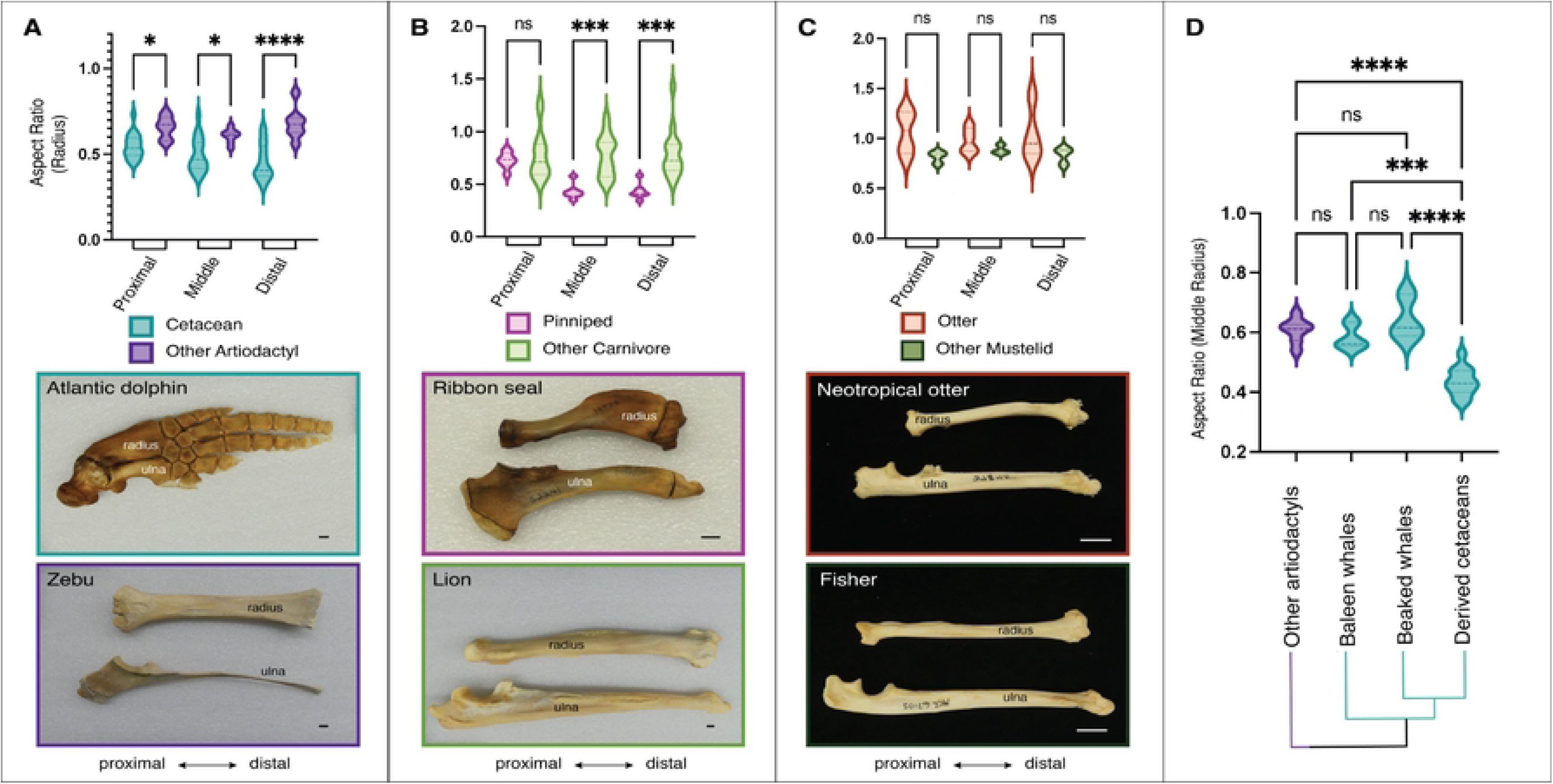
Flattening of the radius convergently emerges in cetaceans and pinnipeds. (A-C) Aspect ratio (AR) along the radial diaphysis and representative photographs for aquatic mammals and outgroups. (A) Cetaceans (n=18 species) vs. other artiodactyls (n=7 species). (B) Pinnipeds (n=7 species) vs. other carnivores (n=14 species). (C) Otters (n=7 species) vs. other mustelids (n=7 species). (D) AR at the middle of the radius for cetacean subgroups. Only derived cetaceans have flattened bones. (Other artiodactyls- n=7 species; baleen whales- n=3 species; beaked whales- n=3 species; derived cetaceans- n=12 species). ns = not significant; * = adj. p value < 0.05; *** = adj. p value < 0.001; **** = adj. p-value <0.0001. Scale bars= 1cm.

Measurements of the radius in pinnipeds (seals and sea lions) and cetaceans (whales and dolphins), two independently evolved clades of diving aquatic mammals with hydrofoil-shaped flippers[11,12], also demonstrated significantly flatter bones compared to outgroups. All pinnipeds except for the walrus had dramatically flattened radii, and as a group, pinniped species had a significantly flatter radius compared to other carnivores at the middle and distal radial diaphysis (adj. p-value proximal > 0.9999; middle = 0.0007; distal = 0.0003) (Fig 2B and S2). As a whole, cetacean species had flatter radii than other artiodactyls, particularly at the distal end of the radial diaphysis (adj. p-value proximal = 0.0190; middle = 0.0161; distal < 0.0001) (Fig 2A and S2). However, closer examination revealed that within cetacea, *Mysticetes*, the earliest branching cetacean clade, which includes all baleen whales, and *Ziphiidae*, the beaked whales[26,27], did not exhibit a flat radius compared to other artiodactyls (adj. p-value non-aquatic artiodactyls vs baleen whales = 0.8986; beaked whales = 0.6964; baleen vs beaked whales = 0.4489). Derived cetaceans had significantly flatter radii than other artiodactyls or the early-branching whales (adj. p-value vs. non-aquatic artiodactyls < 0.0001; baleen whales = 0.0005; beaked whales < 0.0001) (Fig. 2D and S2). This indicates that whereas flat forelimb bones arose in the ancestor of pinnipeds, this trait only arose in more derived cetacean species. Interestingly, otters, which use their webbed hindfeet and tails, rather than their forelimbs, for propulsion[12] and are notably more agile in land locomotion than pinnipeds, did not exhibit a flat radius compared to other mustelids (adj. p-value proximal = 0.2279; middle > 0.9999; distal = 0.3943) (Fig 2C and S2). However, it should also be noted that otters, as well as non-aquatic mustelids, have a twisting morphology of their radial bones that complicates aspect ratio measurements (Fig S3D). Ulnar bones were also measured across mammal samples, however comparisons of ulnar aspect ratio in mammals were confounded by an extreme reduction and occasional fusion of the ulna to the radius in artiodactyls (Fig S3A-C). These results suggest that flat radial bones are widespread in aquatic species that have convergently adapted to diving-based locomotion in both avian and mammalian classes. Intriguingly, in both clades, this phenotype is limited to species that do not retain ancestral land or air locomotion, suggesting parallel patterns of functional constraint.

The degree of flattening of the radius was significantly higher in penguin species compared to either cetacean or pinniped species, particularly at the proximal end of the bone (adj. p-value vs. proximal cetacean = 0.0007; proximal pinniped < 0.0001; middle cetacean = 0.0059; middle pinniped = 0.2319). This difference was even more pronounced in the ulna, where penguin bones were significantly flatter than those of cetaceans at all points along the ulna (adj. p-value proximal < 0.0001; middle < 0.0001; distal = 0.0017) and significantly flatter than pinniped ulnae at both the middle and distal end (adj. p-value proximal = 0.0513; middle and distal < 0.0001) (Fig S4). As penguins display more extreme flattening in their radius and ulna than any other measured taxa, they can serve as a hallmark example of the evolution of this trait.

### Progressive evolution of altered forelimb aspect ratio in penguin lineages

To gain insight into the evolution of bone widening in the penguin lineage, we focused on the penguin ulna. We measured the width of the ulna along its length, comparing ulnar proportions of modern and extinct penguins with those of the chicken, a common lab model, and the northern fulmar, a close relative of the penguin lineage that does not exhibit altered forelimb aspect ratio (Fig 3). Ulnae from chicken and norther fulmar are widest at the ends of the shaft.

**Fig 3.**
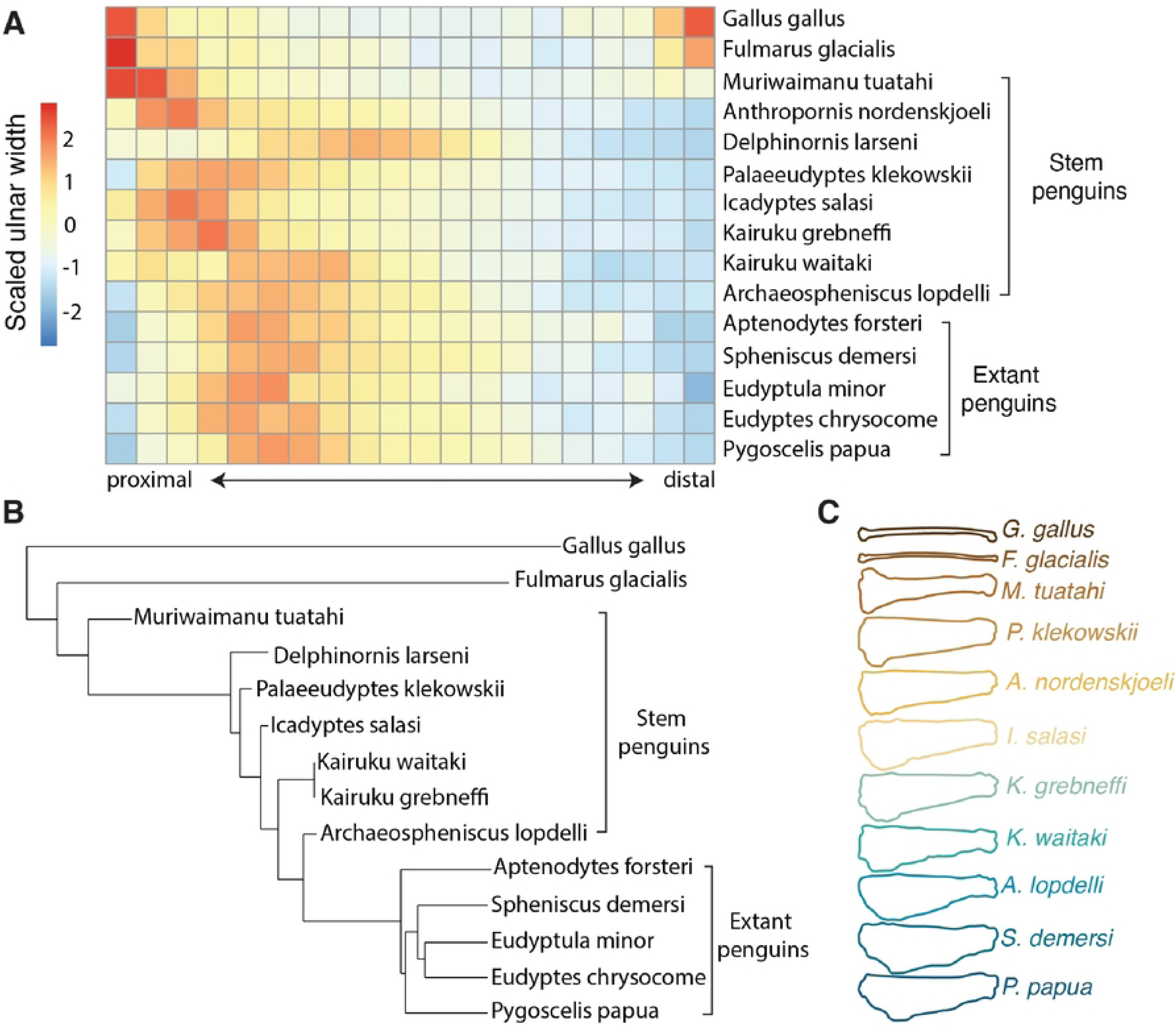
Progressive lateral expansion of the ulna in fossil penguins. (A) Scaled width along the length of the ulna in extant penguins and stem penguin fossils compared to outgroups, chicken (*Gallus gallus)* and Northern fulmar (*Fulmarus glacialis*). Values for each location along the ulnar length represent the number of standard deviations from the mean at that point. (B) Phylogeny of stem and extant penguins (from Cole et al., 2022). (C) Ulnar shape (dorsal view) of extant and extinct penguins.

In contrast, modern penguin ulnae are widened along the length of the central shaft and are widest approximately a third of the length from the proximal end of the bone. In ancestral penguins, we observed a pattern of progressive extension of the olecranon, the bony prominence at the proximal end of the ulna that articulates with the humerus and serves as the attachment point for the triceps tendon. In early stem penguins (*Muriwaimanu tuatahi*[28] and *Anthropornis nordenskjoeli*[29]) the olecranon extends slightly distally into the central shaft of the bone. Later stem penguins (*Palaeeudyptes klekowskii*[30] and *Icadyptes salasi*[31,32]) show an intermediate level of distal olecranon expansion, and in fossils of the latest branching stem penguins measured *(Kairuku grebneffi*, *Kairuku waitaki*[29] and *Archaospheniscus lopdelli*[23]), distal expansion of the olecranon is similar to that observed in modern penguins (Fig 3).

Collectively, with the exception of the early branching *Delphinornis larseni*[29], which shows an unexpectedly distal expansion of the olecranon, we find a pattern of bone widening over the course of penguin evolution that progresses from the proximal end of the ulna towards proportions seen in modern penguins. This suggests that flattened bones in the penguin may have evolved by progressive distal widening.

### Penguin forelimb bone flattening occurs through novel development of a lateral bone ridge

The development of the penguin forelimb has not been addressed since a descriptive analysis of embryonic forelimbs of the gentoo penguin (*Pygoscelis papua)* conducted during the second French Antarctic Expedition in 1908-1910[33]. Early descriptive reports[33] showed that penguin-specific forelimb morphologies form *in ovo,* though the timing of the dorso-ventral flattening of the forelimb bones and associated reduction in musculature remain unknown. In principle, such changes in morphology could be the result of evolved alterations to early patterning of the musculoskeletal system in the limb or due to modifications of growth, cell death, or other tissue remodeling in post-patterning/late development stages.

We collected a time course of gentoo penguin embryos (*Pygoscelis papua*) to interrogate the developmental mechanisms underlying the changes in forelimb morphology (Fig 4A,B). Gentoo penguins live in colonies and lay eggs during a relatively synchronized period in mid-late spring, allowing targeted collection of specific embryonic stages. They generally re-lay eggs if lost due to predation or sample collection[34,35], minimizing the ecological impact of harvesting eggs for study.

**Fig 4.**
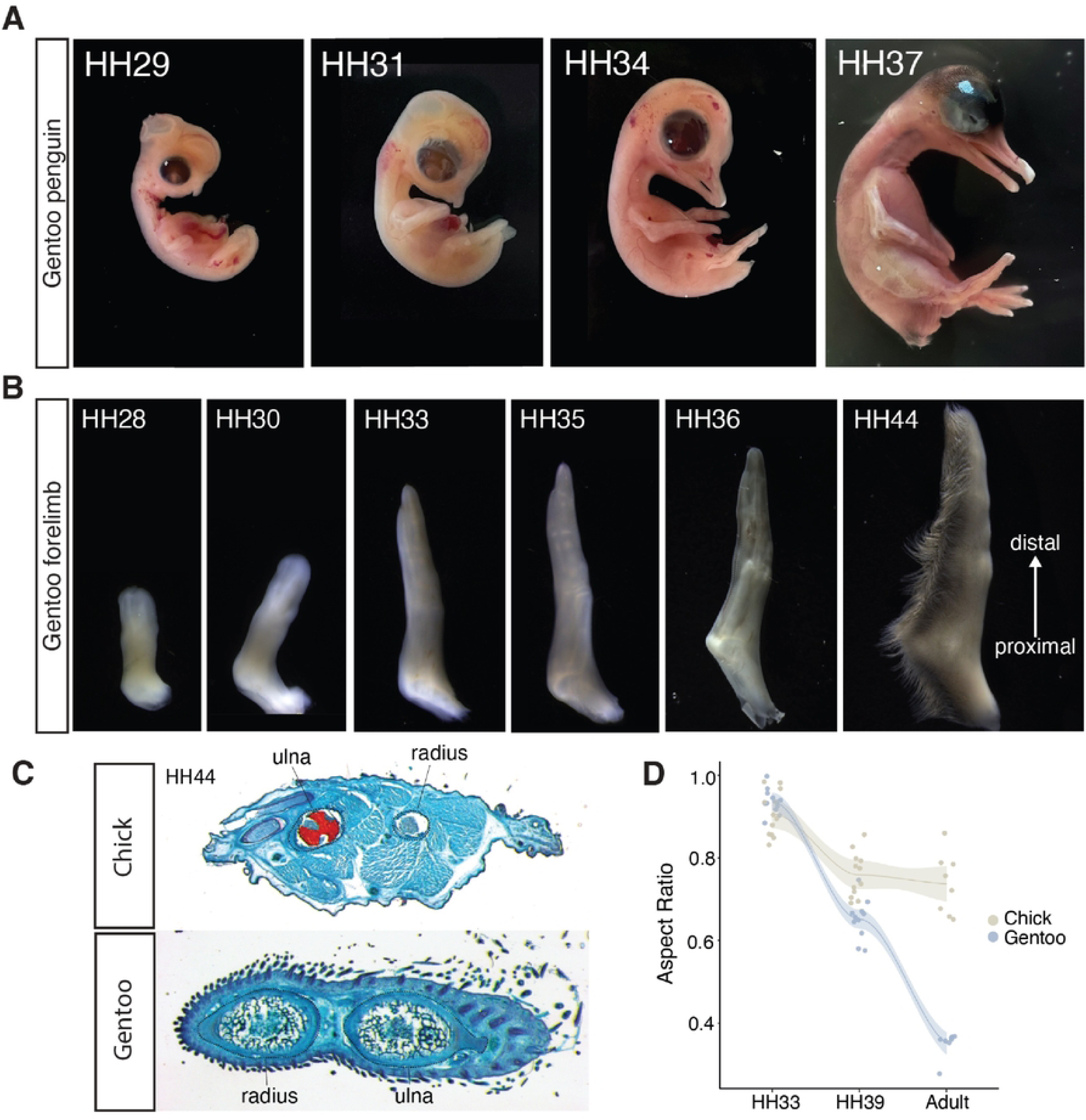
Forelimb bone flattening in the penguin occurs during late embryonic development. (A) Representative images of gentoo penguin embryonic staging (B) Embryonic penguin forelimb development (C) Safranin-O and Fast-green staining of transverse sections through HH44 chick and penguin forelimb showing reduced musculature and altered bone morphology in the penguin (D) Radial and ulnar aspect ratio in embryonic and adult chicken and penguin forelimbs.

We find that gentoo forelimbs in the early embryo (up to HH29-31) largely resemble those of chicks. This similarity changes, however, as the forelimb develops, as they acquire a distinctly flipper-like morphology *in ovo* by HH33 (Fig 4B). Despite the apparent flattening of the forelimb bud as a whole, the mean aspect ratio of the radius and ulna is not significantly different from the chick at HH33 (p-value =0.11), suggesting that the developmental mechanisms that result in limb bud flattening are distinct from those that result in bone flattening. By HH39, the penguin radius and ulna are significantly flatter that the chick (p-value < 6.9e-5), becoming increasingly flatter into adulthood (Fig 4C,D). Thus, the initial morphogenesis of penguin long bones produces rod-like ruminants, round in cross section and similar to that of the chick. It is remodeling during later phases of development that underlies the flattened bone phenotype.

To gain insight into the late differential remodeling in penguins, we performed Safranin-O/Fast Green staining of planar sections of chick and penguin forelimbs at HH38, highlighting the cartilage tissue. Interestingly, at this stage, many gentoo forelimb bones, including the ulna, radius, carpometacarpus 2, and several phalanges have cartilaginous ridge-like structures starting at the epiphyses (the ends of the long bones) and extending down the length of the bone. These ridges are absent in homologous, comparably staged bones of the chick forelimb (Fig 5A). We examined the ulna at several time points in order to understand how these structures change over development. The ulnar ridge emerges at the proximal end of the ulna and progressively extends distally along the lateral edge of the ulna during development (Fig 5B), mirroring the pattern observed in the penguin fossil record. We hypothesize that this lateral ridge results in an anterior-posterior widening of the ulna without commensurate change in the dorsal/ventral “height” of the bone, producing the flattened aspect ratio we observe in adult penguin bones. In concordance with this hypothesis, the distal end of carpometacarpus 2 shows a clear epiphyseal ridge structure, whereas neither end of carpometacarpus 3 has a detectable epiphyseal ridge; in the adult penguin, carpometacarpus 2 has a significantly flattened aspect ratio and is nearly twice as wide as carpometacarpus 3, which remains round (Fig 5A,C).

**Fig 5.**
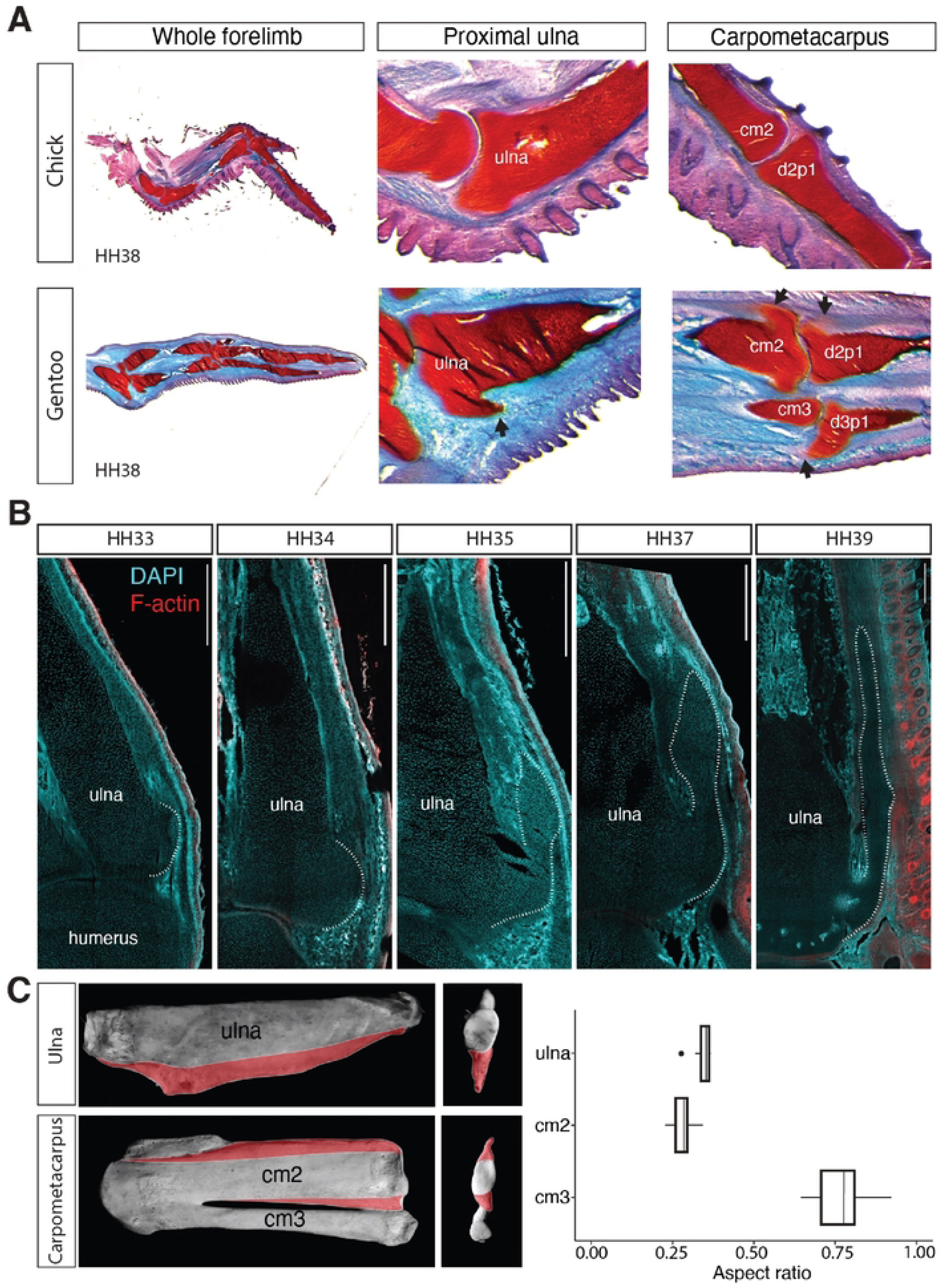
Penguin forelimb bones have bone ridge-like structures. (A) Safranin-O and Fast green stains of planar sections through HH38 chick and penguin forelimbs. Black arrows indicate bone ridge-like structures in penguin bones. (B) DAPI and F-actin in planar sections through the proximal gentoo ulna at embryonic stages from HH33-HH39. The dotted-lines outline the bone ridge-like structure. Scale bars (top right) = 500um. (C) Dorsal and distal views of adult gentoo penguin forelimb bones. Flattened flanges are highlighted in red. Aspect ratio measurements (right) show flattening in ulna and carpometacarpus 2 (cm2) but not carpometacarpus 3 (cm3).

### Muscle loss in the distal penguin flipper coincides with bone ridge expansion

The ridges along the lateral edges of embryonic forelimb bones in the developing penguin resemble bony growths that develop at tendon attachment points in other skeletal elements. In at least one such limb tendon attachment site studied in the mouse, the deltoid tuberosity, mechanical forces exerted by muscle contraction have been shown to be necessary for extended maintenance and growth[36]. We therefore wondered if muscle contraction contributes to the expansion of the penguin ridge structure. Notably, previous descriptions of adult penguin morphology demonstrated a marked lack of musculature in the distal penguin flipper[14]. This does not, however, rule out a role for muscle activity in skeletal shape change in the penguin forelimb, as muscle tissue could, in principle, be transiently present in the developing flipper at the time of ridge formation and regress at later timepoints.

We first asked whether penguin musculature exhibits normal patterning and migration in the early stages of limb development. In vertebrates, muscle progenitors are specified in the ventrolateral edge of the somitic dermomyotome, and then delaminate and migrate into the limb bud, forming dorsal and ventral masses where they proliferate, and differentiate into myoblasts. These muscle masses are subsequently divided into individual muscle bundles which are patterned by lateral-plate mesoderm derived connective tissue cells[37–41]. At HH26, we observed no difference in the pattern of muscle migration between chick and penguin and were able to detect expression of HGF, an early muscle migration cue, TCF4, a marker of the pattern-generating muscle connective tissue, and PAX3/7, muscle precursor markers, in both chick and penguin forelimb (Fig S5A,B).

Given the apparently normal patterning of muscle in the penguin forelimb, we hypothesized that muscle must be lost at later developmental stages. In the chick forelimb, muscle masses in the chick forelimb zeugopod progressively expand between HH33 and HH39 (Fig 6A,E) and show high levels of cell proliferation, measured by phospho-H3S10 (Fig 6D,F). While muscle masses in the gentoo forelimb zeugopod initially have similar numbers of cells as chick at HH33 (Fig 6B,E), they show a significant decrease in muscle cell proliferation over time and do not increase the number of cells per muscle bundle (Fig 6D-F). The decrease in cell proliferation in gentoo muscle cells is not accompanied by an increase in apoptosis (Fig 6C and S5C).

**Fig 6.**
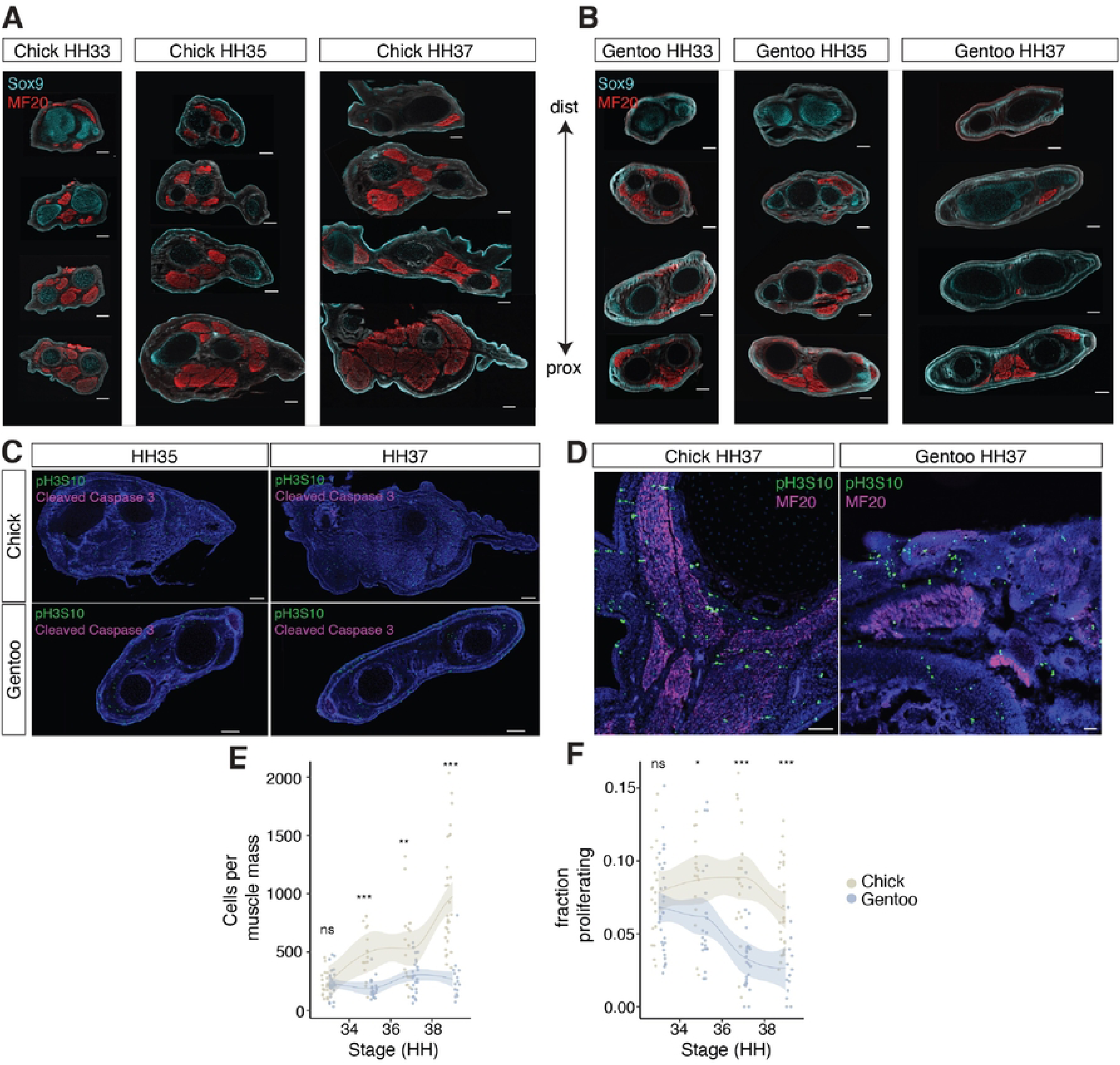
Reduced proliferation contributes to the reduction in gentoo forelimb muscles. (A, B) Immunofluorescence staining of Sox9 and muscle myosin (MF20) in embryonic chick (A) and penguin (B) forelimb. Each column has sections from a single forelimb zeugopod ranging from distal (top) to proximal (bottom). (C) Proliferation (pH3S10) and cell death (cleaved Caspase 3) in chick and penguin forelimb zeugopod. (D) Proliferation and muscle myosin in chick and penguin zeugopod. (E) Number of cells per muscle bundle in chick and penguin zeugopod from HH34-38 (F) Fraction of proliferating muscle cells during development in chick and penguin forelimb zeugopod. Scale bars = 50 um.

Collectively, these results demonstrate that the absence of muscle in the penguin flipper is the result of altered cell proliferation late in development, rather than changes to early muscle specification or migration. The decrease in muscle cell proliferation, which is significant as early as HH35 is largely coincident with the formation of the penguin forelimb’s bone ridges. The relative absence of muscle at these time points therefore makes it unlikely that distal muscle contraction is required for bone ridge maintenance or extension.

### Forelimb bone ridges co-express cartilage and connective tissue markers

If the ridge-like structures observed in the penguin forelimb form through an analogous process as the bone ridges that form at tendon-attachment sites, we would expect similar patterns of gene expression between these tissues. These structures have been best characterized in mouse forelimb, where bone ridge progenitors of tuberosities have been shown to co-express markers of both tendon and cartilage, such as the tendon markers *tenascin C* (TNC) and *scleraxis* (SCX)[42], and the chondrocyte marker *SRY-box 9* (SOX9)[36]. We performed spatially barcoded RNA-sequencing[43] (Light-seq) (Fig 7A) to determine whether the structures we observe in the penguin forelimb are transcriptionally similar to the bone ridges formed at tendon attachment sites in the mouse. Using this targeted technique of genome wide transcriptional analysis, we compared the transcriptional profile of ulnar chondrocytes, ulnar ridge tip cells, and periosteal connective tissue running along the length of the ulna in HH38 gentoo forelimbs (Fig 7A-D). The resulting transcriptomes segregate by tissue of origin in a PCA plot (Fig 7B) and show broad sets of significant differentially expressed genes (DEGs) in each tissue (Fig 7C,D and S6).

**Fig 7.**
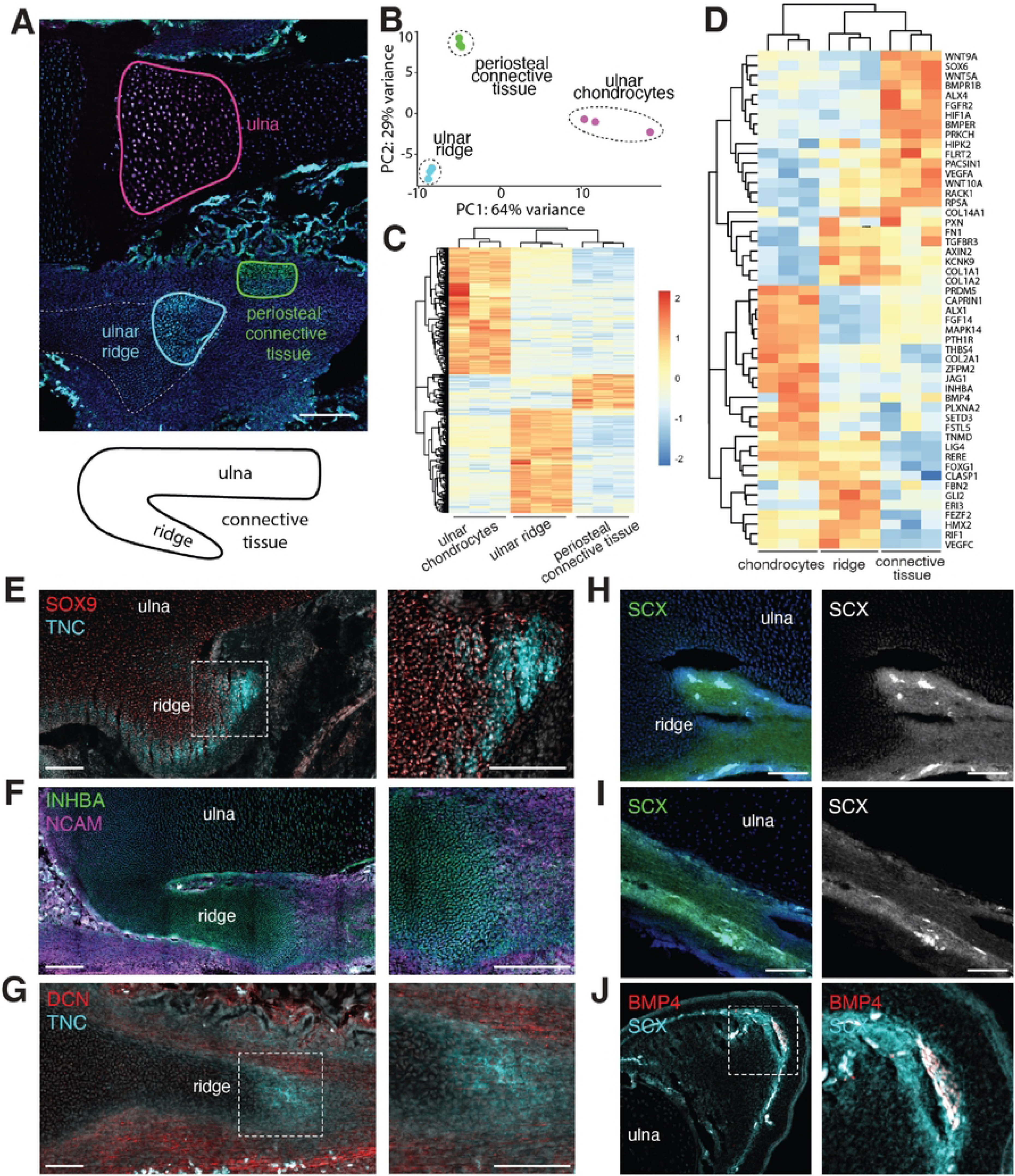
Molecular characterization of penguin bone ridge-like structures. (A) Light-seq spatially barcoded RNA-sequencing of HH38 penguin chondrocytes, ulnar ridge, and periosteal connective tissue. (B) PCA of samples separates by tissue type. (C) Heatmap of all differentially expressed genes in Light-seq data. (D) Heatmap of differentially expressed genes involved in developmental processes. (E) Expression of SOX9 and Tenascin-C (TNC) in ulnar ridge at HH36. (F) Inhibin-A (INHBA) and NCAM expression in the ulnar ridge and surrounding connective tissue at HH36. (G) DCN enrichment in the connective tissue surrounding the TNC-positive ulnar ridge at HH38. (H, I) SCX expression in the tip of the ulnar ridge and periosteal connective tissue at HH36. (J) SCX and BMP4 are co-expressed in the lateral ulna in transverse section. Scale bars = 200uM (A); 100um (all others).

As expected, periosteal connective tissue sampling detected genes known to be upregulated in tendons and ligaments including *fibrillin-2* (FBN2)[44] and *fibronectin-1* (FN1)[45], while ulnar chondrocytes expressed known cartilage markers such as *bone morphogenic protein receptor 1B* (BMPR1B), and *SRY-box 6* (Sox6)[46]. Cells in the newly forming ridge co-express chondrogenic markers such as *bone morphogenic protein 4* (BMP4)[47] and *parathyroid type 1 receptor* (PTH1R)[48], as well as tendon markers such as *tenomodulin* (TNMD)[42]. This is consistent with penguin ulnar ridge cells having an identity similar to that of other bone ridges and tuberosities described in mammals. In addition, penguin ulnar ridge cells express members of the TGFß signaling pathway, which is known in mouse to be required for bone ridge development[36]. In particular, *inhibin A* (INHBA), a subunit of the inhibin and activin complexes that antagonistically signal through TGFß receptors[49], is dramatically upregulated in the developing ridge, while both ulnar cartilage and periosteal connective tissue express the TGFß receptor, TGFßR3 (Fig 7D and S6D). *Activin A receptor type I* (ACVR1), which forms a repressive complex with INHBA that blocks receptor binding[49], was not significantly differentially expressed in our data set but is expressed in all three tissues (Fig S6D and S7B,E).

To visualize expression patterns of bone ridge-related genes in the penguin, we performed antibody stains for markers of cartilage and tendon. In transverse sections at HH38, TNC is enriched on the lateral edges of the radius and ulna (Fig S6E) and in planar sections, TNC is enriched at the tip of the ulnar ridge where it is co-expressed with SOX9 (Fig 7E,G). Consistent with our RNA-Seq results, not all tendon markers are expressed in the developing bone ridge. Both *decorin* (DCN) and *neural cell adhesion molecule* (NCAM) are broadly expressed in the connective tissue running down the length of the ulna but absent from the ridge (Fig 7F,G). We also analyzed expression of *Scx*, a gene which plays a crucial role in tendon development, but which is absent from our RNA-Seq data (most likely due to incomplete annotation of this gene in the gentoo penguin genome). Using cross-hybridizing chicken fluorescent *in situ* probes, we observe that at HH36, when the bone ridge has only minimally elongated, *Scx* expression is present both in the periosteal connective tissue (Fig 7I) as well as in the tip of the ulnar ridge (Fig 7H). In transverse sections at this stage, *Scx* and the cartilage marker, *Bmp4,* are co-expressed on the lateral edge of the ulna where ridges will form, suggesting these factors presage the formation of the bony ridge (Fig 7J). Later, at HH39, *Scx* is broadly expressed throughout connective tissue as well as the bone ridge but, as expected, is largely depleted in ulnar chondrocytes (Fig S7A,D).

Several genes in our database were expressed at particularly high levels in the bone ridge suggesting activity during ridge extension. *Rere (Arginine-Glutamic Acid Dipeptide Repeats)*, a transcriptional repressor that plays multiple roles in development[50], is expressed broadly in the forelimb but is notably more prevalent in the bone ridge and connective tissue than the ulnar chondrocytes at HH39. Interestingly, levels of *Rere* appear to be much stronger in the connective tissue near the proximal origin of the bone ridge (Fig S7C) than at the developing tip (Fig S7F), suggesting that connective tissue gene expression may differ along the length of the bone ridge. At HH36, INHBA protein is present in all three tissues, but strongly upregulated at the tip of the bone ridge (Fig 7F). At HH39, we find that its partner, *Activin A receptor type I* (ACVR1) is expressed most strongly in the connective tissue, but with substantial expression throughout the developing ridge (Fig S7B,E).

Thus, lateral bone ridges in the penguin ulna express markers of the bony eminences that form at legitimate tendon attachment sites, including genes previously identified as essential for formation of tuberosities such as members of the TGFß signaling pathway.

### Developmental programs associated with penguin bone ridges show evolutionary signatures of selection

Having identified gene sets associated with the developing ulnar ridge in penguins some of the pathways expressed in these tissues, we capitalized on available genomic datasets to ask whether these genes were under selective pressure in flipper-bearing lineages. To that end, we asked if our differentially expressed genes (DEGs) were enriched in two previously published datasets: one of genes with signatures of selection in penguin genomes[51] and another of genes convergently evolving in marine mammals[52]. We found that high-confidence DEGs (False Positive Rate: q < E-10) are significantly enriched in the dataset of genes under selection at the ancestral branch of penguins (p = 0.040), indicating that these genes are associated with the evolution of penguin-specific adaptations. When we distinguish between tissue comparisons, genes that are differentially expressed in the bone ridge compared to the ulnar cartilage are significantly enriched for selection in penguin (p = 0.049), whereas the genes found to be differentially expressed in the ridge compared to local connective tissue are not (p=.991) (Fig 8C and Table S2).

**Fig 8.**
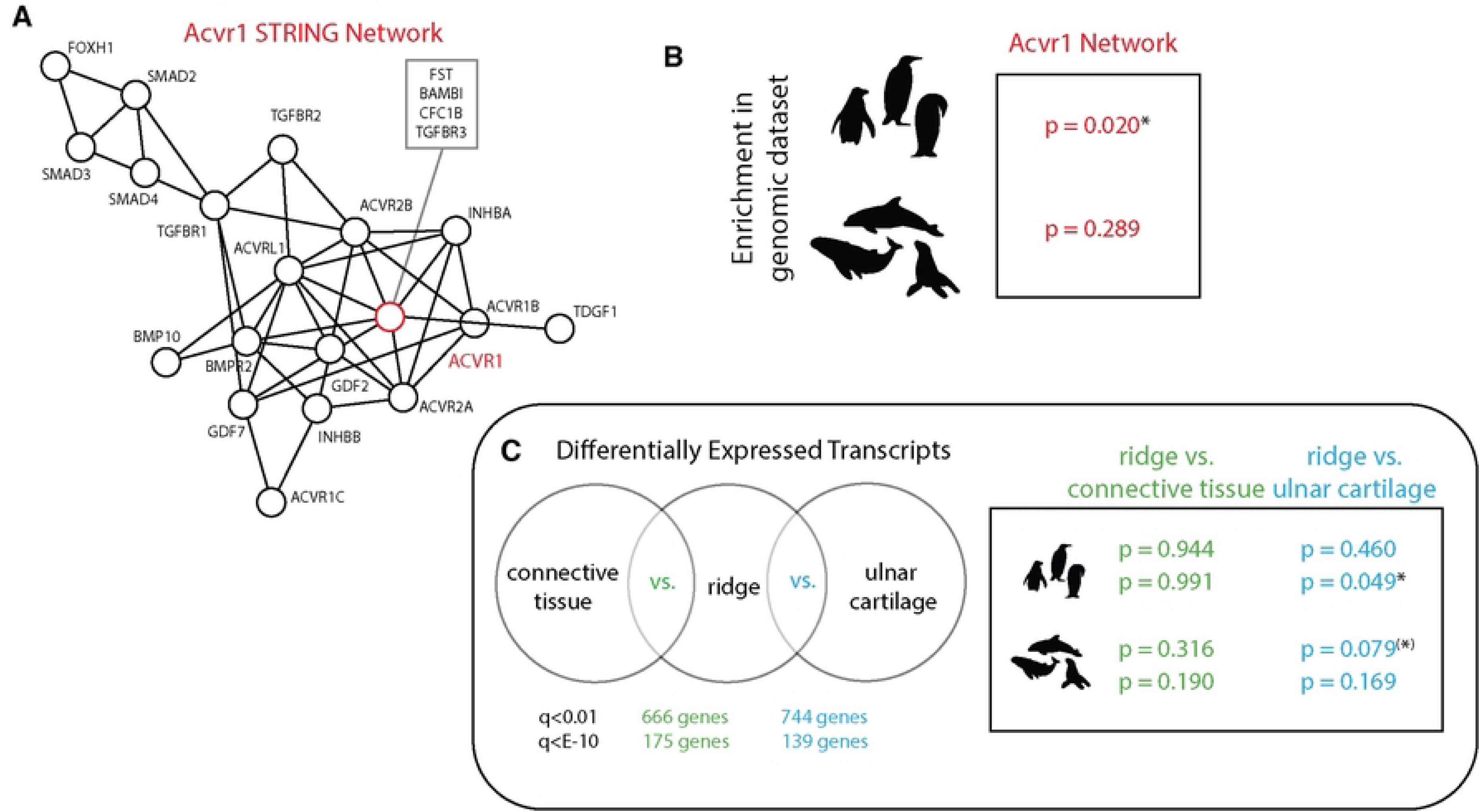
Bone ridge DEGs and ACVR1 interactors are enriched for selection in penguins. (A) ACVR1-interactor network. (B) ACVR1-interactors are significantly enriched in a dataset of genes under selection in penguins but not marine mammals. (C) Differentially expressed genes (DEGs) between the ulnar ridge vs. connective tissue (green) and vs. cartilage (blue) at two stringency thresholds (q < 0.01, top; q < E-10, bottom). High confidence ridge vs. cartilage DEGS are significantly enriched in a dataset of genes under selection in penguins. A less stringent set of ridge vs. cartilage DEGs are nearly significantly enriched in a dataset of convergently evolving genes in marine mammals. * = p < 0.05; ^(^*^)^ = p < 0.10.

This association is maintained when we examine the marine mammal convergence dataset, but only when including a broader and less stringent set of DEGs (q < 0.01), achieving significance at a more modest threshold (p < 0.10). Once again, ridge vs. cartilage DEGs were significantly enriched in the marine mammalian dataset (p = 0.079), whereas ridge vs. tendon DEGs were not (p = 0.316) (Fig 8C and Table S2).

A particularly interesting genetic signal that emerged from our differential expression data in the penguin was inhibin/activin signaling, showing a dramatic increase in expression of *Inhba* in the developing bone ridge as well as expression of *Acvr1* throughout the ridge and strongly in the surrounding connective tissue. Inhba/Acvr1 signal through the TGFß pathway to regulate bone and tendon formation, and this pathway is specifically required for bone ridge formation in mouse[36]. Mutations in the kinase domain of *Acvr1* have also been shown to cause progressive bone over-growth in the human disorder, Fibrodysplasia Ossificans Progressiva (FOP)[53]. Members of the inhibin/activin network were also previously documented to be rapidly evolving in secondarily marine mammals. For example, in the marine mammal convergence dataset, diving mammals have differential sequence evolution in *Acvr2b* (p = 0.0325)[51]. We also identified a number of penguin-specific mutations in the ligand-binding domain of *Acvr1* (Fig S8), however, a computational assessment of these mutations did not predict a dramatic impact on protein function. To determine whether the inhibin/activin network was under selective pressure in the penguin and marine mammal lineages, we developed a STRING network based on Acvr1 interactions (Fig 8A). We tested the members of this network for enrichment in the penguin and marine mammal datasets. Members of the Acvr1 interaction network were not enriched in the marine mammal convergence data set (p = 0.289), however they were significantly enriched for selection in ancestral penguins (p = 0.020) (Fig 8B and Table S2), suggesting that this pathway may have contributed to penguin specific-adaption.

## Discussion

The highly derived morphology of the flipper is a fascinating example of specialization of the vertebrate limb. The observed integration of dramatic shifts in morphology across multiple tissue types - including tendon, muscle, and bone - has allowed a diverse array of vertebrate lineages to adapt to aquatic environments. In the penguin, flat bones, in concert with muscle loss, tendonization, and an expansion of sesamoid bones, likely reinforce the rigidity of the forelimb, allowing it to function more effectively as a stiff flipper for wing-powered diving. The altered flat form additionally allows the flipper to function as a hydrofoil, providing lift during underwater ‘flight’[11,12,14–18,20]. These highly derived features are not unique to the penguin, having emerged convergently multiple times in aquatic tetrapods.

In this work, we find that in birds and mammals, flattened forelimb bones convergently evolved in multiple independent aquatic lineages whose underwater locomotion relies on the use of hydrofoil-like flippers. Intriguingly, we *only* observe forelimb bone flattening in birds and mammals that have lost alternate forms of forelimb-based locomotion. Penguins, mancalline auks, and great auks, birds that have distinctly flattened bones, all lost the ability to fly. In contrast, diving birds that retain flight, including wing-powered divers like petrels, do not exhibit flat bones. This trend extends to pinnipeds which have massively diminished their ability to walk and run (instead using their abdominal muscles to undulate across land), as well as cetaceans, which have lost all land-based locomotion. Otters, which, despite being aquatic mammals, do not use their forelimbs as flippers and retain agile land locomotion, do not exhibit flat forelimb bones. Collectively, this pattern may be due to functional constraint such that flat limb bones are adaptive in species that use their forelimbs as hydrofoils but are extremely maladaptive in the context of non-aquatic locomotion, such that their evolution is only possible in animals that no longer require their forelimbs for their ancestral locomotion. Consistent with this hypothesis, fossil penguins only developed forelimb skeletal modifications after the presumed loss of flight around the K-T boundary[23,52].

Penguins, whose forelimb bones have the most extremely modified aspect ratio of any species we measured, develop externally, *in ovo*, making them suited for developmental analysis of the development of these highly derived limb phenotypes. Through histology and spatial transcriptomics, we demonstrate that the flattening of the forelimb skeleton is not due to alterations to the dorsal-ventral axis, or “height” of the endochondral bones as they first emerge as cartilage models. Instead, we describe a novel bone ridge-like structure that forms subsequently, starting at the epiphysis of multiple long bones in the penguin forelimb and gradually extending along the length of the bones during late embryogenesis, effectively expanding the “width” of the skeletal element. Bones that have such ridges, including the ulna, radius, and carpometacarpus 2, are significantly flattened in the adult, whereas bones that lack this structure, like carpometacarpus 3, remain round. This suggests that the presence of the embryonic ridge is tightly correlated with, and may be largely responsible for, the flat bone phenotype in the adult. While the penguin is the only species we could investigate embryonically, the presence of a sharp ridge-like structure running along the lateral edge of the great auk radius suggests that a similar mechanism of bone ridge expansion may have convergently evolved in that species as well (Fig S1G). The pinniped radius and ulna also superficially appear to have structures resembling bone ridges which extend part of the way along the diaphysis, potentially pointing to a broader convergence of this mechanism of altering bone aspect ratio. Other highly specialized ridge-like structures have evolved in avian hindlimbs such as the large, crested patella in extinct diving bird *Hesperornis*[54] and the rotular process (or cnemial crest) in loons (Fig S1K)[55], both of which are kick powered divers. As such, the expansion of bone ridges may represent a common mechanism supporting the evolution of specialized vertebrate locomotion.

Our analysis shows that, in penguins, the developing bone ridges remain separated from the long bone cartilage by connective tissue through at least HH39, just a few days before hatching. We therefore hypothesize that bone remodeling to facilitate fusion of each long bone with its associated ridges into a single contiguous element occurs extremely late in embryonic development, or perhaps postnatally. Interestingly, the development of ridges along the forelimb bones during embryogenesis parallels the gradual expansion of ulnar width observed in ancestral penguin fossils. We therefore posit a model for the evolution of progressively flatter penguin forelimb bones wherein more derived taxa increase the duration and extent of bone ridge extension and ossification into the periosteal connective tissue, before the final fusion into a single cohesive element occurs.

To probe the mechanism by which lateral ridges along penguin forelimb bones develop, we examined gene expression in this tissue through spatial transcriptomics. We find that the bone ridges that form along the penguin ulna co-express markers found in tendon, connective tissue, and bone. Expression of many of these genes mirrors patterns observed during the normal development of tendon attachment sites (entheses) in rodent models, and also in human pathologies that involve ectopic tendon ossification at the enthesis, including ankylosing spondylitis[56] and the formation of bone spurs[57,58]. Cells at the tip of the developing penguin bone ridge co-express *Scx* and BMP4 in a pattern similar to the mouse deltoid tuberosity.

Initiation of deltoid tuberosity formation is driven by SCX-dependent BMP signaling from the nascent tendon, while its growth and maintenance depends on mechanical forces transmitted from the muscle by the tendon[36]. TNC, a tendon marker[42], which is dramatically upregulated in the tip of the developing penguin bone ridge, has been implicated in new bone formation at the enthesis in ankylosing spondylitis[56]. TGFß activity, which is modified in the penguin bone ridge through the altered expression of both ligands and receptors, is both necessary for the development of normal entheses (tendon-bone interfaces)[36] and sufficient to induce enthesopathies in mouse models when ectopically expressed[59]. In particular, *Inhba*, which inhibits TGFß signaling, is strongly upregulated throughout the bone ridge. Although activins, the signals that promote TGFß signaling in opposition to *Inhba*, were not differentially expressed in our dataset, we identified several penguin-specific mutations in the ligand binding domain of the activin receptor, *Acvr1*. This was particularly interesting because hypermorphic mutations in the kinase domain of *Acvr1* result in ectopic bone growth in humans[53]. Together, these patterns suggest similarities in the mechanisms underlying bone ridge formation at tendon attachment sites in developing mice, the modified penguin flipper, and human pathologies.

Entheses have been shown to be extremely plastic in response to mechanical force, during both embryonic development and in adult animals, to allow adaptation of the force transferring structure[9,60]. In fact, some entheses have been shown to be dependent on transfer of force from muscle contraction to achieve their adult morphology. For example, although initiation of the deltoid tuberosity in mice does not require muscle activity, continued growth is dependent on mechanical input from muscle contraction[36]. Adult penguins have extremely diminished musculature in the distal forelimb[14], raising the possibility that the extension of bone ridges in the forelimb could be muscle-independent. Here, we detailed a decrease in penguin embryonic musculature that is largely contemporaneous with ridge formation. The coincident timing of muscle loss and bone ridge extension makes it unlikely, but not impossible, that penguins rely on distal muscle contraction for bone ridge extension and maintenance.

Surprisingly, we found that the loss of forelimb muscle in the penguin is achieved not through reduction in early muscle precursor migration, nor through muscle cell death by apoptosis, but rather through a reduction in embryonic muscle proliferation in late embryogenesis. This finding mirrors reports that muscle loss in the distal jerboa hindlimb occurs despite normal muscle precursor migration and shows no increase in apoptosis or necrosis[61,62]. The persistence of normal myoblast migration into the distal limb in dramatically different taxa that ultimately exhibit distal muscle loss could suggest a degree of developmental constraint on early muscle patterning in development, possibly because disrupting early steps of muscle formation could have pleiotropic maladaptive effects.

Within the ulnar chondrocytes and connective tissue, we identified differentially expressed gene sets (DEGs) that have signatures of selection in the penguin lineage, suggesting that modifications to some of these genes may have contributed to the evolution of the flipper. In particular, members of an Acvr1 developmental network and bone ridge DEGs were enriched in a dataset of genes under selection in the penguin. Intriguingly, although the bone ridge co-expresses cartilage and connective tissue markers, only genes that were differentially expressed in the ridge vs. cartilage comparison were enriched for selection in penguins. This suggests that non-cartilage genes expressed in the bone ridge (genes exclusive to the bone ridge as well as genes shared between bone ridge and connective tissue) are under stronger selection and thus may be driving the evolution of flattened bones.

It is possible that mammalian flippers convergently evolved through parallel developmental mechanisms, perhaps even involving modifications to some of the same genes under selection in the penguin. Indeed, although Acvr1 interactors were not strongly enriched for convergent evolution in marine mammals, we did see mild significance when we looked at ridge vs. cartilage DEGs from the penguin. Although this finding is constrained to a selection of DEGs that is less stringent on inclusion to the gene set, the enrichment of these genes in the aquatic mammal dataset was very near the typical cutoff for significance (p = 0.079). Previous work, describing convergently evolved genomic elements in marine mammals, supported our findings here, detailing enrichment for tendon-associated genes[51]. The finding of any pattern of shared mechanistic association between clades that convergently evolved these highly multigenic limb modifications, despite some 300 million years of divergent evolution, is quite remarkable. This raises the tantalizing possibility that there may be true convergent overlap in the mechanism(s) that produce forelimb specializations in aquatic mammals and birds.

Together, we find that the muscular and skeletal morphology of the penguin flipper does not arise from changes in the early patterning of the limb, but rather from post-patterning modifications of existing structures during late embryonic development. The flattening of penguin forelimb bones is established through deployment of a common developmental program that also produces bone ridges at tendon attachment sites. We suggest that over the course of penguin evolution, these bone ridges may have extended further and further along the length of the bone, producing the proportions observed in modern penguins. Although embryonic investigation of other species with flipper-like morphologies remains inaccessible, we demonstrate parallels in phylogenetic shifts of bone structure and genomic signals of selection in birds and mammals that use their forelimbs exclusively for aquatic locomotion. Collectively, these results represent an important step towards understanding the evolution and development of highly specialized limb morphologies.

## Materials and Methods

### Sample collection

All embryos were collected in accordance with the appropriate Institutional Animal Care and Use Committee (IACUC) guidelines. White leghorn chicken embryos were obtained from Charles River (MA) and gentoo penguin embryos were collected from Weddell Island in the Western Falkland Islands under Falkland Island research permit number R29/2022. Chicken embryos were incubated at 38°C until they reached appropriate stages and gentoo eggs were collected on targeted dates to obtain specific stages. Embryonic chick tissue was dissected in PBS and fixed in 4% formaldehyde and processed for embedding in OCT and sectioning. Gentoo tissue was dissected in PBS, and stored and shipped in 4% formaldehyde and stored at 4°C until processed for embedding in OCT for sectioning.

### Immunofluorescence, HCR in situ hybridizations, and histology

Fixed embryos were dehydrated through a sucrose gradient to 37% sucrose and embedded in 50:50 ratio of 37% sucrose:OCT (Tissue-Tek). Sectioning was performed on a Leica CM3000 cryostat. For immunohistochemistry, sections were incubated with primary antibodies in PBTS (PBS/BSA 0.2%, Triton 0.1% / SDS 0.02%) overnight, washed 2×10 minutes in PBST, incubated for 1 hour with secondary antibodies washed 2×10 minutes in PBST, and counterstained with DAPI. Primary antibodies used were anti-SOX9 (Millipore), anti-phosphoH3S10 (Millipore), anti-Cleaved Caspase 3 (Cell Signaling), anti-TNC (Millipore), INHBA/Activin A (Invitrogen), NCAM (Invitrogen), and DCN (Abcam) at 1:500 dilutions and MF20, Pax3, and Pax7 (all Developmental Studies Hybridoma Bank) at 1:50 dilutions. Custom HCR in situ hybridization probes for chicken *Scx* and *Bmp4* were ordered from Molecular Instruments (MI). Custom HCR probes for penguin *Acvr1, Rere,* and *Pth1r* were designed in house and ordered as oligo pools from Integrated DNA Technologies (IDT).

HCR RNA-FISH was performed by washing OCT from slides in 1X PBS 2 x 5 min and immersing slides in 70% EtOH for 1-2 hours. Slides were washed twice in 1X PBS for 5 min at room temperature and incubated with pre-heated hybridization buffer (Molecular Instruments) to 10 min at 37C. Slide were then incubated in pre-heated hybridization buffer with final concentration of 40-60nM for MI probes and 10nM for IDT probes at 37C overnight with a coverslip in a humidified chamber. Slides were then washed 2 X 30 minutes with probe wash buffer (MI) then 2 X 100% 5X SSCT for 20 min at room temperature. The slide was dried and pre-incubated with 250uL amplification buffer (MI) in a humidified chamber for 30 min at room temperature. Hairpins H1 and H2 were prepared by separately snap cooling H1 and H2 at 95C for 90 seconds then cooling 30 minutes at RT in the dark. 4 uL of each hairpin were added to 500uL amplification buffer (MI) to a final concentration of 48 nM and incubated together to 5 min at RT in the dark. Slides were dried and incubated overnight with hairpin solution in a humidified chamber. Slides were next washed 3 X 20 min in 5X SSCT at RT, incubated with DAPI and mounted in Fluoromount-G for imaging.

Safranin-O/Fast green stain was performed by washing cyosectioned slides in ddH2O for 10 minutes, staining with 0.08% Fast green for 5 minutes, rinsing in 1.0% Acetic acid to 10 seconds (without rinsing), staining with 0.1% Safranin-O for 3 minutes followed by a single dip into 0.5% Acetic acid, dehydration with 2X washes 80% EtOH and 2X washes in Xylene. Samples were mounted using Permount and imaged on a Nikon dissecting scope.

### Aspect ratio measurements

Adult bone measurements from mammals and birds were performed on specimens in the collection of the Museum of Comparative Zoology (MCZ), Harvard University **©** President and Fellows of Harvard College, using digital calipers (Table S1).

Avian species: At least 2 measurements each of the height (dorso-ventral direction) and width (anterior-posterior) were taken at the proximal, middle, and distal regions of the diaphysis. At least two radii, two ulnae, and two tibiae were measured for each species.

Mammalian species: Single measurements of height (dorso-ventral) and width (anterior-posterior) were taken at three points along the diaphysis. In pinniped species with pronounced ridges, measurements were taken at the widest point of the ridge, just below the ridge, and at the rounded/unmodified end of the diaphysis. Two radii, two ulnae, and tibiae were measured when possible. For the purposes of sub-group comparisons (e.g., cetaceans vs. other artiodactyls), for each species, all measurements at a given point (proximal, middle, distal) were averaged and a single value representing that species was used for statistical testing.

Embryonic specimens: aspect ratio was measured in ImageJ (FIJI) using transverse sections through the middle of the zeugopod stained with DAPI and imaged on a confocal microscope.

Extinct species: Measurements of extinct species were taken from the literature or by tracing and measuring published photographs of radius and ulna[24,25,28–32], except in the case of the great auk, where we measured radius and ulna of samples from the MCZ (**Table S1**).

Relative width of extinct penguins along the length of the ulna was calculated using pheatmap "scale" in R, which subtracts the mean (centering) and divides by the standard deviation (scaling).

Statistical testing: Comparisons between measurements from adult bones were performed in GraphPad Prism using One-Way ANOVA. We corrected for multiple hypothesis testing using a Tukey multiple comparisons test when all categories were compared (**Fig. 2D, S1C, S4**), or a Bonferroni multiple comparisons test when a subset of categories were compared (**Fig. 2A-C**). Comparisons between embryonic aspect ratios were performed in R using Welch’s t-test.

Images of adult radius and ulna and whole articulated great auk skeleton were taken of specimens in the MCZ, Harvard University, **©** President and Fellows of Harvard College (Table S1). Whole articulated chicken skeleton image is from Wikimedia commons (Museum of Veterinary Anatomy FMVZ USP, Wagner Souza e Silva).

### Light-seq

Fixed HH38 gentoo forelimb zeugopods were dissected in 1x PBS and graded through 7% sucrose into a 1:1 solution of OCT (Tissue-Tek) and 30% sucrose in 1x PBS for freezing and cryosectioning at into 20-µm sections on poly-L-lysine coated Ibidi chamber slides. Slides were centrifuged at 800 rcf for 5 minutes to promote adhesion, and antibody retrieval was performed by incubating slides immersed in 10nM sodium citrate pH 6 and incubated in a steamer for 10 min. In situ RT and A-tailing, barcoding, cross-junction synthesis, displacement, and library preparation was performed as described previously^47^ with the following modifications. We used 2 mg/ml sheared salmon sperm DNA (Invitrogen) and 10% dextran sulfate in the barcode hybridization mix to reduce non-specific barcode binding. During the displacement step, we pipetted the RNase H containing solution up and down several times every 15 minutes during the 45-minute incubation at 37°C to increase yield of recovery. Samples were tagmented with a Nextera XT library preparation kit (Illumina) using custom primers for the i5 end as previously described^47^. Sequencing was performed on a NextSeq500 with a read length of 150 and 30% PhiX spike-in. All replicates were pooled together and sequenced across two lanes of a NextSeq500 run.

Sequencing data analysis was performed using published Light-seq analysis code on the Harvard Medical School O2 cluster (Kernel 2.10.0) with Python (v3.7.5), PyTable (v6.1), samtools (v1.12), pysam (v0.17.0), numpy (v1.21.4), pandas (v1.4), Biopython (v1.79), and scikit-bio (v0.5.6). Briefly, Barcode, UMI, and cDNA sequences were extracted from Read 1 using UMI-tools (v1.1.1). cDNA sequence was then mapped to the Pygoscelis papua genome (GenBank assembly accession: GCA_010090195.1). Reads were assigned to genes using FeatureCounts using fractional read counting and the GFF annotation ‘CDS (-M --fraction -g gene_id -t gene) and deduplicated (per gene) with UMI-tools dedup71. Reads were parsed out by barcode sequences using a custom python script^47^. Differential gene expression analysis was performed in R (v6.1) using DESeq^63^. Volcano plots were produced using ggplot2 (v4.0), and enriched genes were plotted using the pheatmap function (v1.0.12).

### Comparative Genomic Analysis

Enrichment of Penguin tissue DEGs, in both the Penguin Ancestral Evolution dataset[52] and the Marine Mammal Convergence dataset[51] was performed using the SUMSTAT method, which has demonstrated power, flexibility, and simplicity over competing gene set enrichment analyses[63–65]. In short, log-transformed p-values for each gene were summed and compared to a distribution of randomly sampled scores of the same set size to determine an enrichment p-value. Any gene IDs which did not map between datasets were dropped. The gene sets at various Q-thresholds are subsets of each other, such that they are not independent tests and multiple hypothesis corrections would be inappropriate.

The phylogenetic trees for mammals in **Fig. 2D, S2, S3A** were generated using PhyloT (https://phylot.biobyte.de) based on NCBI Taxonomic numbers (**Table S1**). The cetaceans were further refined by averaging relative branch length values for baleen and beaked whales from the literature[26,27].

The Acvr1 STRING Network was generated using STRING (https://string-db.org) using Acvr1 from *Homo sapiens*.

## Acknowledgements

We thank Marie Manceau for bringing the adaptation of the penguin wing skeletal elements to our attention and for critical help acquiring permits for penguin egg collection. We thank Jeremiah Trimble, Kate Eldridge, Mark Omura, and Madeleine Mullon at the Harvard Museum of Comparative Zoology for assistance with museum collections; Emma West and Josie Kishi for advice on Light-seq protocols; Marie Manceau, Camille Curantz, and Maria Castro Scherianz for help with gentoo sample collection; Lewis Clifton and the staff of Weddell Island for assistance with sample collection and logistics; Chaitra Prabhakara for generous assistance with HCR while H.A.G. was unable to work with formamide; and Robin Carpenter for assistance with mammal bone measurement. Imaging was performed at the MicRoN core at Harvard Medical School and RNA-sequencing was performed using the Broad Walk-up sequencing service at the Broad Institute. This work was funded by NIH grant HD 032443 to C.J.T. Support for H.A.G. was provided by the Lallage Feazel Wall Fellowship of the Damon Runyon Cancer Research Foundation, DRG-2447-21, and NIH grant K99GM157508. Comparison of gene selection supported in part through NIH grant HD115763 to MPH.

## Author contributions

C.L. and C.J.T. conceived the study. All authors contributed to experimental design. C.L. and H.A.G. collected embryonic penguin and chick samples and performed measurements and photography of extant avian and mammalian skeletons. C.L. performed histology, immunofluorescence, microscopy, image quantification, Light-seq, sequencing analysis, and analysis of fossil penguin bones. H.A.G. performed statistical analysis of adult aspect ratios.

C.L. and H.A.G. performed HCR and related microscopy. S.T. performed genomic and bioinformatic analyses. C.L. and H.A.G. constructed figures. C.L. and C.J.T. wrote the initial manuscript draft and H.A.G., M.P.H., and C.J.T. prepared subsequent drafts with significant input and advice from all authors.

## Supporting Information Captions

**Fig S1. Penguin forelimb bones are flatter than those of other avian species.** (**A-B**) Radius and ulna aspect ratio (AR) for avian species at the proximal (**A**) and distal (**B**) epiphyses. (**C**) Mean AR at the middle of the radius and ulna for each species compared by avian subgroup. a, b, and c are significantly different from one another. (flightless- n=3 species; flying- n=8 species; flying + diving- n=12 species; Mancallinae- n=4 species; stem penguin- n=5 species; modern penguin- n=5 species). (**D-J**) Radius and ulna of representative birds: (**D**) little penguin, (**E**) king penguin, (**F**) gentoo penguin, (**G**) great auk. Arrows indicate apparent bone ridge along the radius. (**H**) Transverse view of radius and ulna from wandering albatross (flighted, diving) and gentoo penguin (flightless, diving). (**I**) Lateral view of radius and ulna from northern fulmar and (**J**) red-faced cormorant. (**K**) An elaborate bone ridge at the proximal end of the tibia forms the cnemial crest in the common loon.

**Fig S2. Flattening of the radius in aquatic mammal species by phylogeny.** Aspect ratio measurements near the middle of the radius for mammal species. Colors indicate mammalian clades.

**Fig S3. Complications of measuring aspect ratio of forelimb bones in mammals**. (**A**) Mean aspect ratio measurements near the middle of the ulna for individual mammal species. (**B**) Total fusion of radius and ulna in llama makes measurements of ulnal aspect ratio impossible. Dotted line indicates region of fusion. (**C**) Dramatic reduction and partial fusion of radius and ulna in water buffalo and other bovids complicates measurements. (**D**) The otter radius (left) twists, such that they are flattened at the distal end in the anterior-posterior plane, rather than the dorsal-ventral plane. The otter ulna (right) is easily measured. Scale bars = 1cm.

**Fig S4. Penguin forelimb bones are flatter than those of aquatic mammals.** Mean aspect ratio measurement for penguin (n=5), cetacean (n=18), and pinniped (n=7) species at three points along the diaphysis of the radius or ulna. ns = not significant; **adj. p-value < 0.01; ***adj. p-value < 0.001; ****adj. p-value <0.0001.

**Fig S5. Early muscle migration into the forelimb is not detectably altered in gentoo penguin.** (**A**) Muscle migration into the chick and penguin forelimb at HH26. (Scale bars, 200um). (**B**) Early muscle patterning genes HGF, TCF, PAX7, and PAX3 are detectable in HH26 penguin forelimbs. (Scale bars, 500um). (**C**) Positive control for Cleaved Caspase 3 staining in lateral edge of digit 3 in penguin hindlimb. (Scale bar, 50um)

**Fig S6. Differentially expressed genes from spatial Light-seq of penguin forelimb.** Volcano plot of significantly upregulated genes in HH38 chondrocyte vs periosteal connective tissue (**A**) and chondrocyte vs ulnar ridge (**B**). (**C**) Expression of FGF receptors, ligands, and Notch pathway genes (**D**) Expression of developmental genes of interest from Light-seq data. (**E**) Quantification of Tenascin-C (TNC) localization in transverse sections through HH38 gentoo penguin zeugopods.

**Fig S7. Spatial expression of *Scx*, *Acvr1*, and *Rere* at HH39**. Fluorescent in situ hybridization of penguin ulnar ridge at HH39. (**A, D**) *Scx* transcripts are largely confined to the connective tissue and ridge and are more highly expressed at the tip of the developing ridge than at the junction with the ulna. (**B, E**) *Acvr1* is most highly expressed in connective tissue, but is also abundant in the developing ridge. (**C, F**) *Rere* is expressed in all tissues at similar levels near the tip of the developing ridge, but is more highly expressed in connective tissue near the junction with the ulna.

**Fig S8. Penguin-specific mutations in Acvr1**. Protein alignment of Acvr1’s ligand binding domain colored by similar chemical properties and conservation (predicted by Clustal X). The red star marks the base of the penguin lineage. Red arrows at the bottom indicate penguin-specific mutations identified in Acvr1.

**Table S1.** Accession numbers for extant and fossil species measured.

**Table S2:** Bone ridge DEGs and ACVR1 interactors are enriched for selection in penguins and marine mammals.

## Notes

### Competing Interest Statement

The authors have declared no competing interest.

